# *Enterococcus faecalis* persists and replicates intracellularly within neutrophils

**DOI:** 10.1101/2025.07.03.662968

**Authors:** Claudia J Stocks, Ronni AG da Silva, Haris Antypas, Navin Jeyabalan, Siu Ling Wong, Kimberly A Kline

## Abstract

Chronic wound infection is a major global public health issue, with *Enterococcus faecalis* among the most commonly isolated pathogens from such wounds. Neutrophils are short-lived immune cells critical for host defence, yet *E. faecalis*–neutrophil interactions are poorly understood. Here, we show that instead of eliminating *E. faecalis*, neutrophils provide a niche for intracellular persistence and replication, potentially prolonging infection and inflammation at the wound site. In murine wound beds and ex vivo wound cells, intracellular *E. faecalis* was detected in recruited neutrophils at 24 h post-infection (h p.i). Unexpectedly, extended infection did not induce neutrophil death. Rather, *E. faecalis* infection significantly prolonged the lifespans of both murine and human neutrophils *in vitro* compared to uninfected controls. Quantification of intracellular CFU revealed that *E. faecalis* were phagocytosed regardless of opsonisation and persisted intracellularly through to 24 h p.i. This finding was confirmed via transmission electron microscopy and confocal microscopy. Blinded quantification and fluorescent D-amino acid staining, which marks newly synthesised bacterial peptidoglycan, revealed active replication within murine neutrophils between 6-18 h p.i., followed by a predominately persistent phase between 18-24 h p.i. Infected murine neutrophils remained immunologically active, secreting pro-inflammatory and chemoattractant cytokines. These findings highlight an underappreciated intracellular lifestyle for *E. faecalis* that may underly its ability to persist in chronic wounds and contribute to biofilm-associated infections.

## Introduction

Chronic wound infection represents a major global public health concern, impacting both healthcare costs and patient quality of life [1, 2]. *Enterococcus faecalis* is a Gram-positive opportunistic pathogen associated with infections in a range of contexts including urinary tract infection, endocarditis, neonatal sepsis, and chronic wound infection [3–7]. A facultative anaerobic bacteria and commensal of the human gastrointestinal tract, *E. faecalis* exhibits both intrinsic and acquired antibiotic resistance [8], making these infections inherently and increasingly difficult to treat. Long-considered an extracellular pathogen, *E. faecalis* aggregates and forms biofilms, enhancing its persistence capacity in chronic infections [9]. However, increasing evidence across a range of host cell types point to an intracellular niche and lifestyle of this multifaceted persistent bacterium [10–13].

Neutrophils are short-lived, highly antimicrobial cells of the innate immune system that are the first cells to be recruited to sites of infection [14]. Bacterial phagocytosis typically triggers neutrophil death (apoptosis) [15–18], however some intracellular pathogens can subvert this process to persist or proliferate within neutrophils [19–22]. Research on the interactions between *E. faecalis* and neutrophils is scarce. Existing studies, all >25 years old, focussed on early stages of the interaction (10–120 min), including attachment [23], phagocytosis [24, 25], and rapid killing [26–28]. These reports identified the importance of serum opsonisation for effective neutrophil killing of *E. faecalis* [26, 27]. In the absence of serum opsonisation, substantial uptake (but not killing) of *E. faecalis* expressing aggregation substance was reported in neutrophils [24, 28] and macrophages [29]. Overall, these studies have led to the assumption that neutrophils effectively eliminate *E. faecalis*.

In a mouse model of wound infection, we previously observed *E. faecalis* microcolonies in the wound bed together with a pro-inflammatory cytokine response and an influx of polymorphonuclear leukocytes [30], similar to that observed with cutaneous *Staphylococcus aureus* [31] and *Pseudomonas aeruginosa* infections [32]. Despite this robust acute inflammatory response, *E. faecalis* can persist in murine wounds for at least 7 days [30]. This discrepancy between *in vitro* evidence of rapid neutrophil clearance and long-term *in vivo* persistence presents a knowledge gap in our understanding of *E. faecalis*-neutrophil interactions [7].

Here, we investigated the neutrophil response to *E. faecalis* following the observation of clusters of *E. faecalis* within neutrophils in murine wound tissue at 24 h post-infection (p.i.). Examining neutrophil-*E. faecalis* interactions *in vitro*, we unexpectedly found that phagocytosed *E. faecalis* infection suppressed, rather than induced, neutrophil death in both mouse and human cells out to 24 h. Transmission electron microscopy revealed *E. faecalis* residing within spacious vacuoles inside neutrophils. Intracellular quantification and use of a fluorescent dye to mark newly formed bacterial cell walls revealed that *E. faecalis* was not only persisting, but replicating within murine neutrophils. Multiplex cytokine analysis of *E. faecalis*-infected murine neutrophils identified the release pro-inflammatory cytokines to signal for macrophage recruitment. These findings challenge the prevailing view of neutrophils as fully effective against *E. faecalis*, and further our growing appreciation of its capacity for intracellular replication.

## Results

### *E. faecalis* intracellular clusters are present within neutrophils during wound infection

To investigate the mechanisms underlying *E. faecalis* persistence in wounds, we visualised and charactered its localisation *in vivo* using our established mouse wound infection model [30]. Wounds were inoculated with 10^6^ CFU *E. faecalis* OG1RF for 24 h, then cryosectioned and immunostained for Streptococcal Group D Antigen to detect *E. faecalis* before imaging via confocal microscopy. Tile scanning revealed widespread *E. faecalis* clusters throughout the wound bed **(Fig 1A)**, including both extracellular microcolonies and apparent intracellular clusters **(Fig 1B-C)**. This observation is consistent with previous reports of viable *E. faecalis* within both CD45^+^ (immune) and CD45^-^ (non-immune) wound cell populations at 1-3 days p.i. [12].

**Figure 1:**
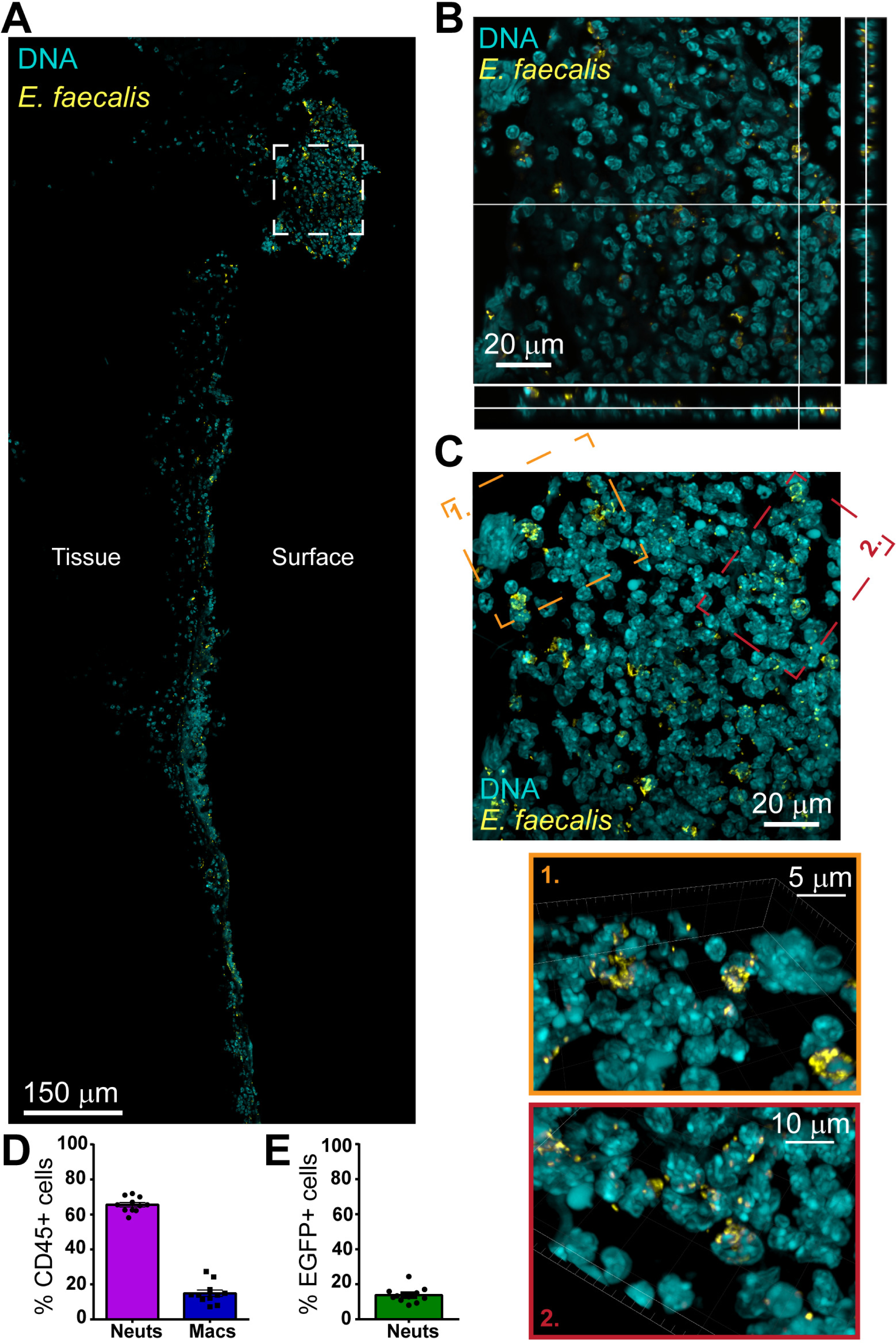
Intracellular clusters of *E. faecalis* within neutrophils in the murine wound bed at 24 h p.i. Mice were wounded and infected with 10^6^ CFU of *E. faecalis* OG1RF **(A-C)** or OG1RF pDasher **(D, E)** for 24 h. **(A-C)** Wounds were excised, fixed and embedded before being cryosectioned, mounted on slides and stained for Streptococcal Group D antigen to mark *E. faecalis* bacteria (yellow) and DAPI for DNA (cyan). Sections were imaged at 63x on a confocal microscope. **(A)** Depicts a tile scan of a portion of the entire wound bed, with tissue and surface sides indicated. White inset box displayed as **(B)** a single slice from the z stack with ortho view and **(C)** max intensity projection, with additional zoomed in angled orange and red insets of IMARIS 3D reconstructions. **(A-C)** Are from a single experiment and mouse, representative of N=6 mice from 3 independent experiments. **(D-E)** Wounds were excised, then cell dissociated from this tissue via liberase reagent. Cells were then stained and imaged via a BD LSRFortessa X-20 Cell Analyzer. **(D)** Displays the relative proportion of (Ly6G^high^, CD11b^+^) neutrophils compared to (Ly6G^low^, F4/80^+^, CD11b^+^ macrophages in the CD45+ immune cells analysed. **(E)** Further displays the proportion of neutrophils that were EGFP (i.e. *E. faecalis* OG1RF pDasher positive). **(D, E)** Depict mean + SEM from N=6 mice, from 3 independent experiments.

To determine host cell types containing intracellular *E. faecalis*, we repeated the wound infection using a GFP-fluorescent strain of OG1RF, *E. faecalis* pDasher [33]. Single cells were isolated from infected wounds and stained for immune cell markers (CD45^+^) to discriminate between neutrophils (Ly6G^high^, CD11b^+^) and macrophages (Ly6G^low^, F4/80^+^, CD11b^+^) **(Supplementary Fig 1A)**. Of the immune cells detected, ∼70% were neutrophils, and ∼15% were macrophages **(Fig 1D)**. Among the neutrophils, ∼15% were GFP^+^, indicating the presence of *E. faecalis* pDasher **(Fig 1E, Supplementary Fig. 1B)**. Given the likely pDasher plasmid loss during infection, this is probably an underestimation. These findings demonstrate that *E. faecalis* clusters reside intracellularly within neutrophils during wound infection.

### *E. faecalis* suppresses neutrophil death

Given the presence of intracellular *E. faecalis* within neutrophils in the murine wound bed, alongside our knowledge of *E. faecalis* persisting long term in these wound beds [30], we hypothesised that they may evade neutrophil killing to persist intracellularly. To examine this, we first assessed the impact of *E. faecalis* exposure on neutrophil viability. Primary neutrophils were isolated from murine bone marrow and exposed to *E. faecalis* OG1RF or *Staphylococcus aureus* USA300 (multiplicity of infection [MOI] 1) for 6 h. Cell death was assessed by via lactate dehydrogenase (LDH) release into the culture supernatant. As expected, *S. aureus* infection increased cell death, consistent with previous reports that phagocytosis accelerates neutrophil apoptosis [18, 34]. Unexpectedly, *E. faecalis* infection resulted in negligible or even negative (reported here as zero) cell death **(Fig 2A)**, likely due to bacterial interference with LDH readings [35]. To correct for this, we used a modified LDH assay that couples measurements from both the supernatant and the cell lysate, allowing estimation of the surviving cells based on retained LDH **(Fig 2B)**. This revealed ∼90% survival of *E. faecalis*-infected neutrophils at 6 h p.i., significantly more than 70% for the uninfected control or ∼60% following *S. aureus* infection **(Fig 2A)**. Extending the infection for 18-24 h did not increase cell death beyond that of uninfected control cells **(Supplementary Fig 2A, B)**. Immunofluorescence microscopy supported these findings, showing *E. faecalis*-infected neutrophils still attached to the coverslip, with intact multi-lobulated nuclei and minimal morphological signs of activation **(Fig 2C)**. By contrast, *S. aureus* infected neutrophils exhibited extensive cell death, with nuclear condensation, and cells destroyed or overrun with bacteria **(Fig 2D)**. Together these data suggest that *E. faecalis* infection delays or inhibits neutrophil death typically triggered by infection [18].

**Figure 2:**
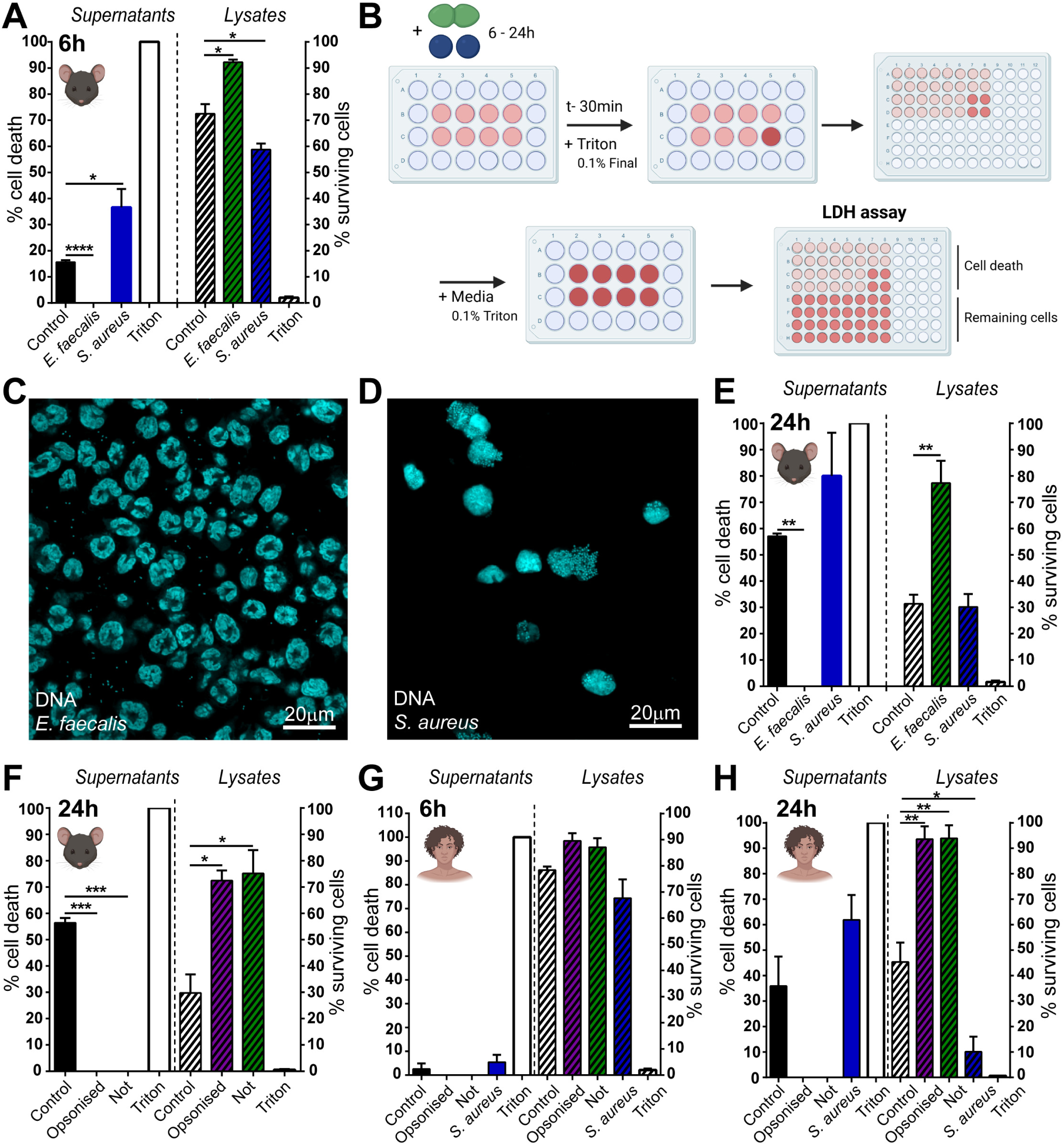
Suppression of spontaneous neutrophil cell death by *E. faecalis* infection. Neutrophils were infected with *E. faecalis* OG1RF or *S. aureus* USA300 (MOI 1) for 6 h, followed by gentamicin exclusion of extracellular bacteria. **(A, E-H)** At the indicated timepoint, LDH assay was performed both on supernatants and remaining cells lysed identically in media. Total percentage cell death in both cases was calculated by comparison to 100% kill well ‘Triton’. **(B)** Schematic diagram depicting the modified LDH assay which determines both percentage cell death and percentage surviving cells for the given timepoint. Light pink depicts cell culture media during standard assay, dark pink once Triton has been added to lyse all remaining cells. Experiment is performed on cells in a 24 well plate before media is taken and assay performed in a 96 well plate. **(A, C-F)** Depict experiments with primary murine neutrophils, **(G, H)** with human. For **(F-H)** *E. faecalis* were additionally incubated prior to infection for 15 min in either 10% mouse **(F)** or human **(G, H)** serum ‘Opsonised’ or else 10% FBS ‘Not’. **(C, D)** At 6 h p.i., cells were fixed with 4% PFA and stained with propidium iodide to mark DNA, then imaged at 63x on a confocal microscope. Data **(A, E-H)** depict (n=6, 4, 4, 3,3) experiments, respectively, mean+SEM; Data was analysed using One Way ANOVA with Dunnett’s multiple comparisons test. * denotes p<0.05, ** p<0.01, ***p<0.001, ****p<0.0001. **(C, D)** are representative of n=4 experiments.

To further characterize the fate of infected neutrophils, we added a high dose of gentamicin directly to the media at 6 h p.i. to kill extracellular bacteria, preventing re-infection and enabling monitoring of any ongoing intracellular infection **(Supplementary Fig. 2C)**. At 18 h and 24 h p.i., only 30-40% of uninfected control neutrophils remained viable, similar to the spontaneous apoptosis observed in the absence of antibiotic treatment, whereas ∼90% and ∼75% of *E. faecalis*-infected neutrophils survived at the same timepoints, respectively **(Fig. 2E, Supplementary Fig. 2D)**. Given previous reports that serum opsonisation enhances *E. faecalis* uptake and killing [24, 26, 27], we tested whether opsonisation affected neutrophil survival. Neutrophils were infected with *E. faecalis* opsonised with either 10% homologous mouse serum or treated with 10% heat inactivated fetal bovine serum (FBS) as a non-opsonised control. At 24 h p.i., ∼70% of *E. faecalis*-infected neutrophils remained viable, regardless of opsonisation, in contrast to ∼30% in control cells **(Fig. 2F)**. We next validated this observation in primary human neutrophils infected with *E. faecalis* in the presence of absence of 10% autologous serum. At 6 h p.i., cell death was comparable across all conditions, except for a slight drop in *S. aureus-*infected neutrophils **(Fig. 2G)**. By 24 h p.i., ∼90% of *E. faecalis*-infected neutrophils remained viable, regardless of opsonisation, compared to ∼40% of control cells and ∼10% of *S. aureus*-infected cells **(Fig. 2H)**. Thus, infection with *E. faecalis* suppresses the cell death of both primary murine and human neutrophils, independent of opsonisation.

To determine if this phenomenon is strain-specific, we tested 7 additional *E. faecalis* strains, including vancomycin-resistant strain V583 [36] and clinical wound isolates [30, 37]. All but one wound isolate (Ef_22) suppressed neutrophil death, comparable to OG1RF **(Supplementary Fig. 2E)**. However, LDH readings varied among strains, with some (OG1RF, Ef_33, Ef_40, Ef_49) showing negative, artifactual readings, while others (Ef_22, Ef_23, Ef_46, V583) provided more accurate readings **(Supplementary Fig. 2F).** Despite these differences most *E. faecalis* strains broadly extended neutrophil life span beyond that of uninfected controls.

### *E. faecalis* is engulfed by neutrophils and persists at late timepoints

To understand whether *E. faecalis*-mediated delay of neutrophil death also affects neutrophil function, we assessed bacterial uptake and intracellular persistence. Murine neutrophils were infected with MOI 1 **(Fig. 3A)** or MOI 10 **(Fig. 3B)**, and CFU enumerated from both the supernatant (extracellular) or the triton lysed neutrophils (intracellular) at 2, 6 and 24 h p.i. At 2 h p.i., opsonisation enhanced infection numbers at 2 h p.i., but this advantage dissipated by 6 h p.i. **(Fig. 3A, B)**. Surprisingly, intracellular *E. faecalis* persisted out to 24 h p.i. regardless of opsonisation. Extracellular CFU increased until 6 h p.i., reflecting bacterial replication in the media. However, following gentamicin treatment at 6 h to kill extracellular bacteria, no viable extracellular CFU were detected at 24 h **(Supplementary Fig. 3A)**. Thus, *E. faecalis* are readily engulfed by murine neutrophils and can persist intracellularly at late timepoints. In human neutrophils infected with MOI 1, intracellular CFU were undetectable at 2 h p.i. and rose to ∼10^3^ at 6 h p.i, regardless of opsonisation. By 24 h p.i., only low levels of opsonised intracellular *E. faecalis* were detectable **(Supplementary Fig. 3B)**. By contrast, infection with MOI 10 resulted in detectable intracellular CFU at all timepoints **(Fig. 3C)**. The levels of *E. faecalis* (MOI 1) in murine neutrophils **(Fig. 3A)** was comparable to that in human neutrophils (MOI 10) **(Fig. 3C)**, suggesting that human neutrophils are more effective at clearing intracellular *E. faecalis.* At 24 h p.i., ∼30% of infected human neutrophils (MOI 10) showed signs of cell death **(Supplementary Fig. 3C)**. Thus, *E. faecalis* can persist within viable human neutrophils for up to 24 h.

**Figure 3:**
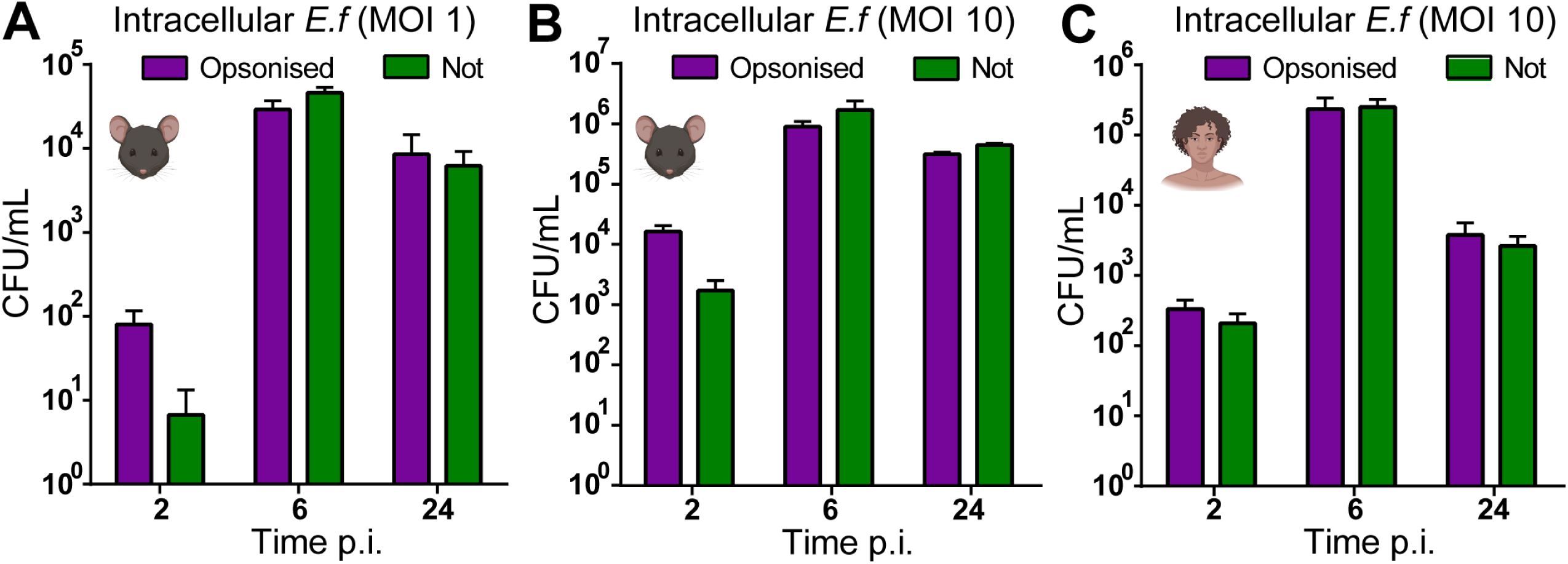
Intracellular persistence of phagocytosed *E. faecalis* in neutrophils to 24 h p.i. Murine **(A-B)** or human **(C)** neutrophils were infected with **(A)** MOI 1 or **(B, C)** MOI 10 *E. faecalis* OG1RF. The infection was halted at either 2 h (for 2 h timepoint) or 6 h, after which extracellular bacteria were killed by gentamicin exclusion. At the given timepoints, wells were gently washed once with PBS, then lysed in PBS + 0.1% triton, serially diluted on BHI for CFU enumeration. Data depicts mean + SEM, from (n=4, 4, 4) independent experiments, respectively.

### *E. faecalis* reside within membrane-bound vacuoles in neutrophils

To gain insight into the intracellular niche for *E. faecalis* within neutrophils, we again infected murine and human neutrophils with *E. faecalis* OG1RF at MOI 1 or MOI 10 respectively, for 6 h, followed by transmission electron microscopy (TEM) **(Figure 4)**. At 6 h p.i., infected neutrophils retained their multi-lobular nuclei and intact plasma membrane, with pseudopodia extending from the cell surface **(Fig. 4A, C, Supplementary Fig. 4A)**. Distinct subcellular structures were visible, including specific (lighter/less electron dense) and azurophilic (darker/more electron dense) granules, glycogen granules, and strands of endoplasmic reticulum (ER). Internalised *E. faecalis* were consistently observed, frequently in large, spacious membrane-bound vacuoles (yellow arrowheads), which were more numerous within human neutrophils (likely reflecting the higher MOI used). In some instances, disrupted vacuolar membranes and visible damaged or degraded bacteria were also observed (magenta arrowheads). Multiple, separate compartments containing one or more bacteria were detected, however they may represent segments of a single larger compartment. At 24 h p.i. **(Fig. 4B, D, Supplementary Fig. 4B)**, infected neutrophils continued to maintain their overall cell morphology, including the nuclear membrane, granule content, and intact ER. In human neutrophils, numerous large, empty vacuoles were evident **(Fig 4C)**, potentially reflecting cleared phagosomes or early apoptotic features. Similarly, several small vacuoles near the plasma membrane were visible in murine neutrophils **(Fig 4B)**. Internalised bacteria in a combination of membrane-bound and membrane-disrupted compartments, were visible in both murine and human neutrophils. Notably, *E. faecalis* within intact membrane-bound vacuoles appeared morphologically healthy, while those exposed to the cytoplasm or associated with granules showed signs of degradation and disintegration. By contrast, control uninfected neutrophils at 24 h p.i. exhibited widespread vacuolisation, nuclear breakdown, and cellular degradation **(Supplementary Fig. 4C, D)**. Uninfected cells that could be visualised at all were overrun with vacuoles and degrading components, with breakdown of the nuclear membrane and nucleus. These striking differences provide ultrastructural confirmation of both the extended neutrophil survival observed in LDH assays **(Fig. 2C, D)** and support the intracellular persistence of *E. faecalis* within both murine and human neutrophils **(Figure 3A, B)**.

**Figure 4:**
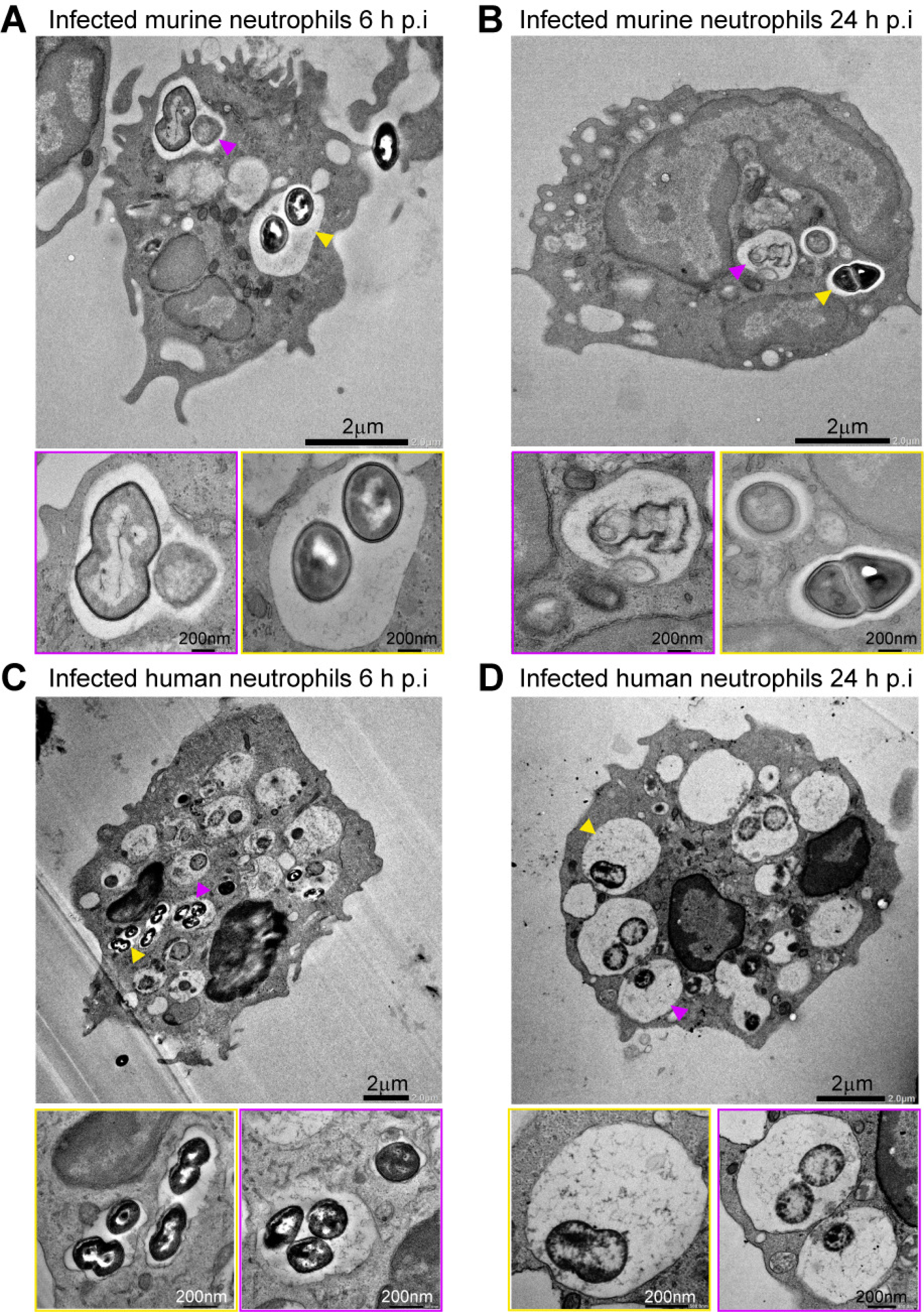
Localisation of *E. faecalis* within membrane-bound vacuoles in murine and human neutrophils. **(A-B)** Murine or **(C-D)** Human neutrophils were infected with **(A-B)** MOI 1 or **(C-D)** MOI 10 of *E. faecalis* OG1RF for 6 h, followed by gentamicin exclusion of extracellular bacteria. At **(A, C)** 6 h or **(B, D)** 24 h p.i., cells were washed with PBS then fixed with 2.5% glutaraldehyde. Cells were imaged via transmission electron microscopy. Depict individual cells from their respective timepoints, representative of at least 6 cells / condition. Yellow arrowheads mark larger, intact membrane bound compartments, with magenta arrowheads a vacuole with a partial/degraded membrane.

### *E. faecalis* replicates intracellularly within neutrophils

To investigate whether *E. faecalis* replicates intracellularly within neutrophils, we stained murine neutrophils with the cytoplasm dye CellTracker Blue, before infection with MOI 1 of *E. faecalis* OG1RF for 6 h. Intracellular bacteria were visible at 6 h p.i. **(Fig. 5A)** and 24 h p.i. **(Fig. 5B)**. Blinded quantification of intracellular bacteria per neutrophil revealed a 3-fold increase between 6 and 18 h p.i., which persisted to 24 h p.i. **(Fig. 5C)**. We also observed a modest, but significant increase in the percentage of infected neutrophils between 6 h and later timepoints **(Fig. 5D)**. Since extracellular bacteria were killed by gentamicin from 6 h p.i., this suggests that the ∼25% cell death observed at 24 h via the LDH assay **(Figure 2D)** predominately comprised uninfected neutrophils.

**Figure 5:**
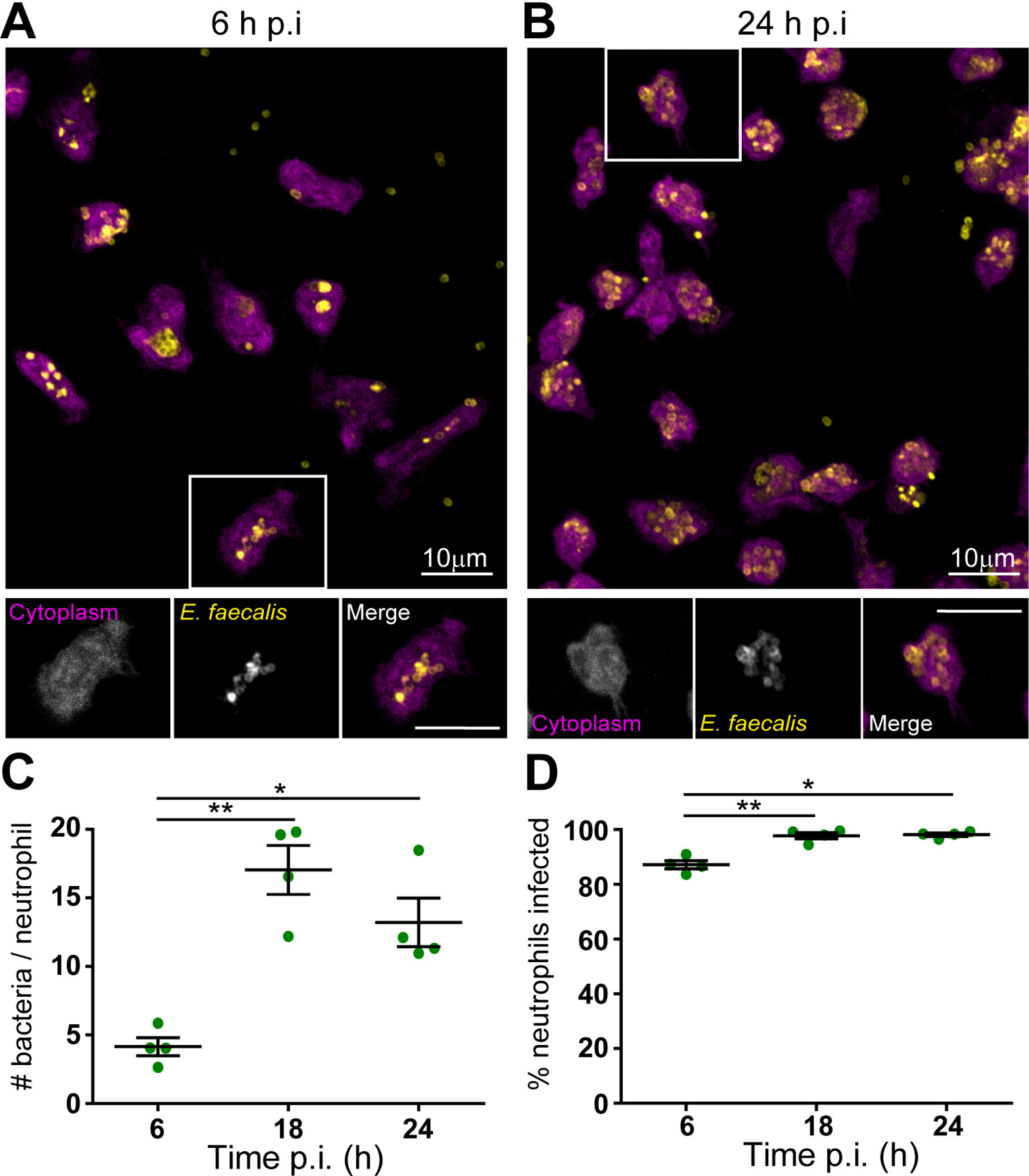
Engulfment and intracellular persistence of *E. faecalis* within murine neutrophils to 24 h p.i. Murine neutrophils were pre-stained with Cell Tracker Blue (cytoplasm, magenta), then infected with *E. faecalis* OG1RF for 6 h, followed by gentamicin exclusion of extracellular bacteria. At **(A, E)** 6 or **(B, F)** 24 h p.i., cells were fixed with 4% PFA and stained for Streptococcal Group D antigen (*E. faecalis* bacteria, yellow) before being imaged at 63x on a confocal microscope. **(C-D, E-F)** Images from (4 per condition, minimum 50 cells total) were blindly quantified to determine the total number of bacteria, neutrophils, and infected neutrophils. **(A-B)** Depict a single experiment, representative of (n=4) independent experiments. **(C-D)** Depict mean ±SEM, from the same (n=4) independent experiments. Data was analysed by One Way ANOVA with Tukey’s multiple comparisons test, * denotes p<0.05, ** p<0.01

To further confirm *E. faecalis* replication within neutrophils, we used the fluorescent D-amino acid probe RADA, which incorporates into newly synthesised bacterial peptidoglycan [38]. Infected murine neutrophils were treated with gentamicin at 6 h p.i., followed by RADA addition at either 7 or 18 h p.i (i.e., 1 or 12 h after gentamicin addition), and imaged at 24 h p.i. When RADA was added at 7 h p.i., we observed fluorescence incorporation in the cell wall of intracellular *E. faecalis*, indicating active replication **(Fig. 6A)**. By contrast, RADA incorporation was minimal when added at 18h p.i. **(Fig. 6B)**, suggesting that *E. faecalis* are replicating intracellularly within murine neutrophils between 6 and 18 h, followed by a persistent non-replicative phase from 18 h onwards.

**Figure 6:** Intracellular replication of *E. faecalis* within murine neutrophils, between 6-18 h p.i. Murine neutrophils were pre-stained with Cell Tracker Blue (cytoplasm, magenta), then infected with *E. faecalis* OG1RF for 6 h, followed by gentamicin exclusion of extracellular bacteria. At **(A)** 7 h or **(B)** 18 h p.i., RADA (fluorescent D-amino acid incorporated into bacterial cell walls during synthesis, cyan) was added into the high dose gentamicin media. At 24 h p.i., cells were fixed with PFA, and stained for Streptococcal Group D antigen (*E. faecalis* bacteria, yellow) before being images at 63x on a confocal microscope. Scale bar in insets depicts 10 μm. Images depict a single experiment, representative of (n=4) independent experiments.

### Infected neutrophils are capable of proinflammatory cytokine signalling

Finally, we sought to determine whether the prolonged intracellular occupation and replication of *E. faecalis* within neutrophils was immunologically ‘silent’ or accompanied by cytokine signalling. Murine neutrophils were infected, left uninfected, or treated with 10 ug/mL lipoteichoic acid (LTA) as a positive stimulus control. Supernatants were collected at 6 and 24 h p.i. and assessed for proinflammatory cytokine release **(Figure 7)**. At 6 h p.i., *E. faecalis*-infected neutrophils secreted modest levels of tumour necrosis factor TNF, granulocyte colony stimulating factor G-CSF, chemokine CXCL1 and Monocyte chemoattractant protein-1 MCP-1. High levels of macrophage inflammatory protein MIP-1α were also detected at levels similar to LTA treated controls. By contrast, secretion of interleukin 6 (IL-6), chemokine RANTES/CCL5, and IL-lβ was induced by LTA, but not by *E. faecalis* at this early timepoint. By 24 h p.i., cytokine production by *E. faecalis*-infected neutrophils was markedly increased. High levels of TNF, IL-6, G-CSF, CXCL1, IL-12 p40, MCP-1, MIP-1α and RANTES were detected **(Figure 7A-H)**. High levels of MIP-1β were also detected at this time, exceeding the limit of accurate quantification by the assay (data not shown). However, in agreement with our data reporting the suppression of neutrophil death by *E. faecalis* infection **(Figure 2)**, minimal IL-1β was detected **(Figure 7I)**. Thus, *E. faecalis-*infected neutrophils have the capacity to signal for immune cell recruitment and activation, in particular other innate immune cells such as macrophages and additional neutrophils.

**Figure 7:**
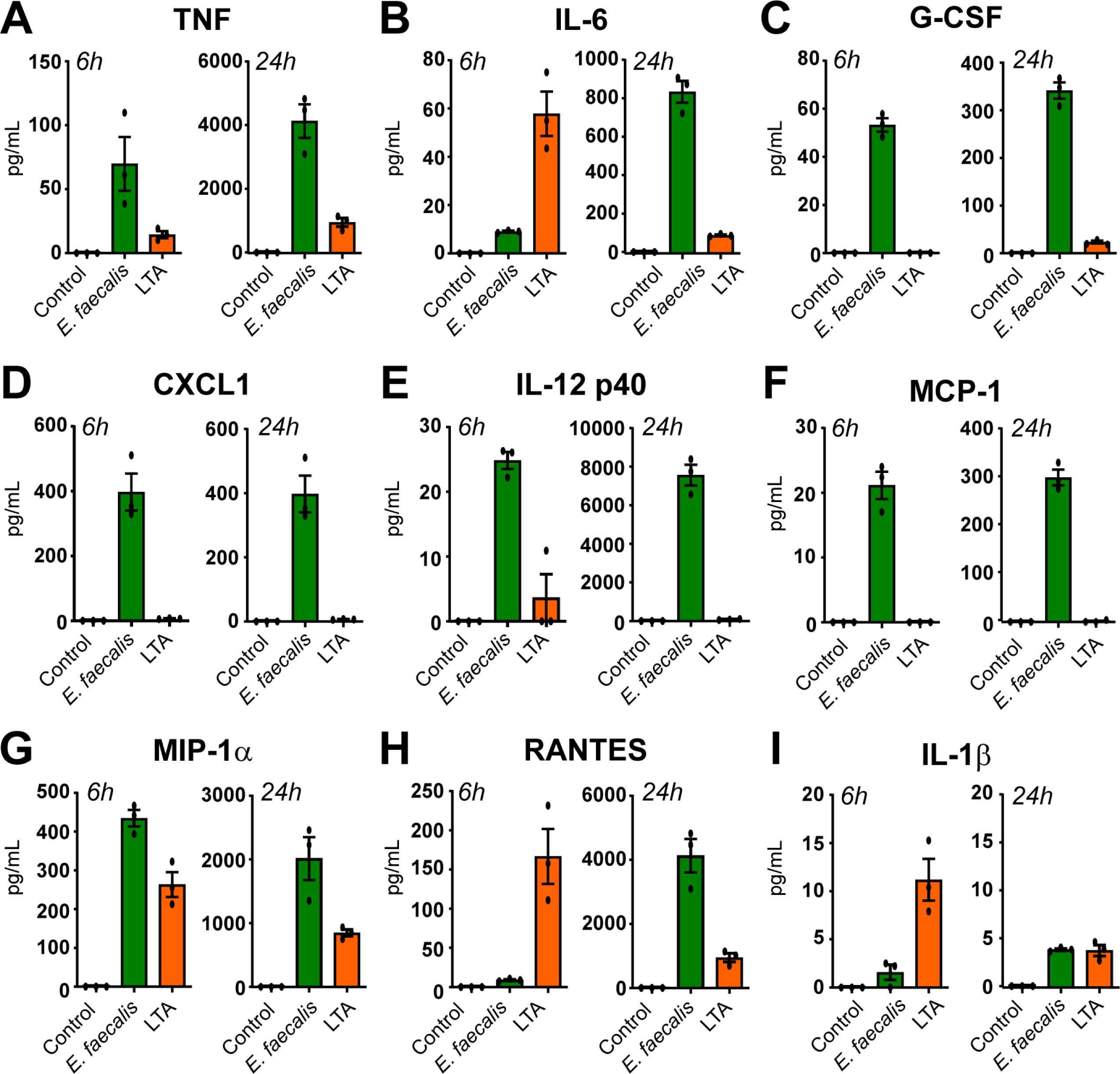
Robust neutrophil cytokine response induced by *E. faecalis* infection. Murine neutrophils were infected with MOI 1 of *E. faecalis* OG1RF or stimulated with 10ng/mL lipoteichoic acid (LTA) for 6 h followed by gentamicin exclusion of extracellular bacteria. At either 6 or 24 h p.i., supernatants were removed and assessed via multiplex cytokine analysis. Data depicts mean±SEM from 3 independent experiments.

## Discussion

Neutrophils are highly reactive, inherently short-lived immune cells. Whilst examples of bacterial pathogens persisting and/or replicating with macrophages are numerous [39], similar examples in neutrophils are exceedingly rare. Indeed, phagocytosis of most bacteria accelerates neutrophil death [40], as reported for *Escherichia coli* [15], *Streptococcus pnemoniae, Streptococcus pyogenes, Mycobacterium tuberculosis, Listeria monocytogenes, Borellia hermsii* [41]. Some bacteria, such as *S. aureus*, induce rapid neutrophil death via necrosis or necroptosis [42].

Only a handful of intracellular pathogens *Chlamydia pneumoniae* [20], *Neisseria gonorrhoeae* [43], *Francisella tularensis* [44], *Leishmania major* [45] and *Anaplasma phagocytophilum* [46] have been reported to delay neutrophil death or exploit neutrophils as a niche for replication. Thus, the finding that *E. faecalis*, a traditionally extracellular, biofilm-associated pathogen, can suppress neutrophil death and persist and replicate inside these immune cells was highly unexpected. This discovery reshapes our understanding of neutrophil function in response *E. faecalis* and helps resolve the long-standing disconnect between *in vitro* studies reporting efficient neutrophil killing and *in vivo* observations of persistent infection despite an acute immune response [30]. Previous research on *E. faecalis*-neutrophil interactions demonstrated bacterial adherence [23], phagocytosis [24], and killing [26–28], but were limited to very early timepoints (≤2 h p.i.). This narrow temporal approach to studying neutrophil-bacterial interactions is commonplace, possibly leading to underestimation of bacterial capacities to resist neutrophil killing. Similarly, phagocytosis has often been taken as the endpoint, without verifying subsequent bacterial killing. For example, research with extraintestinal pathogenic *E. coli* (ExPEC) revealed that most intracellular bacteria remained viable 3.5 h p.i. following phagocytosis by neutrophils [47]. Similarly, neutrophils obtained from the peritoneal cavity of *S. aureus-*infected mice contained viable bacteria capable of establishing infection when transferred into naïve animals [48].

This work expands our evolving understanding of *E. faecalis* beyond being an extracellular pathogen [13]. Indeed, increasing evidence shows that *E. faecalis* can be taken up by and persist within a range of mammalian cells including endothelial cells [49], epithelial cells [50, 51] and macrophages [11, 52, 53]. Recent studies have shown that *E. faecalis* not only persists but also replicates intracellularly within keratinocytes, RAW 264.7 macrophages, and hepatocytes using BrdU or RADA staining following extracellular antibiotic killing [12, 54]. Interestingly, intracellular bacteria isolated from infected keratinocytes were internalised by naïve keratinocytes at a nearly 10-fold higher rate [12], similar to observations of *S. pyogenes* uptake and subsequent reinfection of macrophages [55].

In our study, *E. faecalis*-infected neutrophils were fully capable of pro-inflammatory signalling to drive immune cell recruitment, which is in keeping with previous *in vivo* wound infection findings at 24 h p.i. [30], and stands in contrast with observations at 72 h and 96 h p.i. immune suppression and M2 macrophage polarisation was observed following *E. faecalis* wound infection [56]. By prolonging neutrophil life span, *E. faecalis* may create a temporary intracellular niche that supports its persistence and replication, leading to increased bacterial burden and enhanced inflammation at the wound site. It is likely that a feedback loop exists involving ongoing phagocyte recruitment, phagocytosis, and intracellular persistence, with short-lived neutrophils facilitating subsequent longer term persistence in macrophages [11, 12]. A similar mechanism has been described for *Leishmania major*, an intracellular parasite that exploits neutrophils as a ‘Trojan horse’ to gain entry into host macrophages [45]. *Leishmania* delays neutrophil apoptosis while inducing MIP-1β secretion, enhancing macrophage engulfment of infected neutrophils, within which the internalised bacteria survive and replicate [45]. This process was visualised *in vivo* in a murine model in which dermal macrophages acquired parasites via direct transfer from, or efferocytosis of, infected neutrophils [57].

TEM of infected neutrophils confirmed that despite infection, murine and human neutrophils retained their morphology and multilobulated nuclei, in striking contrast to uninfected control neutrophils. TEM also revealed that *E. faecalis* reside within spacious, membrane-bound vacuolar compartments at 6 and 24h p.i. This observation is consistent with previous ultrastructural studies of human neutrophils infected with closely related *Enterococcus faecium*, which similarly showed bacteria within discrete, membrane-bound compartments, typically surrounded by substantial ‘empty’ space [27]. TEM of ExPEC within neutrophils also revealed the occasional presence of ‘spacious phagosomes’ alongside ‘tight phagosomes’ [47] as are more commonly observed [58]. Interestingly, TEM of neutrophils collected from murine peritoneal lavages 24 h p.i. with a *S. aureus* clinical blood isolate also revealed bacteria within ‘spacious phagosomes’, whilst a mutant lacking the virulence regulator Sar were predominately within ‘tight phagosomes’ [48].

The mechanism by which *E. faecalis* delays neutrophil death and resists antimicrobial clearance to permit persistence and replication remains to be determined. It is well documented that purified LTA [59] and LPS [60] can delay spontaneous neutrophil death, in the case of LTA via Toll-like receptor 2 and pattern recognition receptor CD14 [59]. Similarly, growth factors such as G-CSF [60, 61] and GM-CSF [62, 63] also prolong neutrophil lifespan, largely through signalling pathways that converge on Nf-κB activation [64]. Thus, in the context of *E. faecalis* infection, one possibility is that initial recognition of LTA on phagocytosed *E. faecalis*, followed by robust production of G-CSF by infected neutrophils, promotes neutrophil survival via NF-kB pathways. However, G-CSF signalling can also enhance neutrophil phagocytic and antibacterial activity [65], which is not consistent with the observed persistence of intracellular bacteria in our study. Further, proinflammatory cytokines such as TNF [66] typically drive neutrophils towards apoptosis, although conflicting reports suggest the impact of this cytokine on neutrophil survival may be time-dependent [67]. Adding further complexity, *E. faecalis* can suppress NF-κB signalling in macrophages [68], raising the possibility that similar immune evasion tactics could be active in neutrophils.

Extensive work has been carried out to characterise the mechanisms by which *F. tularensis* suppresses neutrophil function to establish colonisation [22, 44, 69, 70]. *F. tularensis* inhibits the generation of reactive oxygen species (ROS) during neutrophil infection, even in the presence of compounds that stimulate ROS production [44, 69]. *F. tularensis* infection also interferes with apoptotic pathways, blocking the cleavage and activation of caspase-3, caspase-8 and caspase-9 [70]. Finally, *F. tularensis* preserves neutrophil mitochondrial integrity by preventing Bax translocation and Bid processing during infection [22]. These host pathways offer a useful framework for identifying the molecular mechanisms by which *E. faecalis* may similarly subvert neutrophil death and promote intracellular persistence. Detailed characterisation of intracellular ROS levels or impact on apoptotic pathways during *E. faecalis* infection are lacking, however in murine neutrophils, ROS generation has been associated with bacterial killing as early as 4 h p.i. [71].

The deliberate manipulation of immune cell death pathways is a well-established therapeutic approach in oncology [72], and there is growing interest in applying similar approaches to infectious disease [73, 74]. Understanding how *E. faecalis* suppresses neutrophil apoptosis may therefore uncover novel therapeutic targets. Such insights are particularly relevant for tackling chronic wound infection, as well as a broader array of *E. faecalis-*driven pathologies including infective endocarditis and urinary tract infection [1, 2, 68, 75].

## Materials and Methods

### Bacterial strains and culture

All bacterial strains including *Enterococcus faecalis* strain OG1RF [76], *E. faecalis* strain OG1RF pDasher [33], *E. faecalis* strain VR583 [36] and *Staphylococcus aureus* strain USA300LAC [77] were all grown on brain heart infusion (BHI) medium (BD Biosciences) agar (BD Biosciences). Clinical wound swab isolates were all obtained from Tan Tock Seng Hospital, Singapore [30, 37]. For overnight cultures, a single colony from a fresh bacterial streak was inoculated into in a 10 mL static culture, overnight at 37°C. After twice washing with PBS, *E. faecalis* was adjusted to OD600 nm = 0.5 (∼3 ×10^8^ CFU/mL) and *S. aureus* OD600 nm = 1.5 (5 ×10^8^ CFU/mL) to enable MOI calculations (and serially diluted on BHI agar to subsequently confirm).

### Murine model of wound infection

Mouse wound infections were performed as previously described [30]. Briefly, 6-9 week old mice were anesthetised using isoflurane. All hair is then removed from the backs of mice via (electric) shaver followed by hair removal cream (Nair). Following 70% ethanol disinfection, a wound in made using a 6 mm sterile biopsy punch (Integra Lifesciences). 10ul of bacterial inoculum (10^6^ CFU) is added to the wound, after which Tegederm adhesive dressing (3M) was applied. 24 h post-infection, a ∼1cm^2^ square piece of tissue (encompassing the wound) was collected and processed based on downstream experimentation.

For cyro-sectioning, tissues were excised with approximate dimensions of 1.5 cm by 1.5 cm and placed on Whatman filter paper (Sigma-Aldrich). Tissues were immediately submerged in 2mL of 4% paraformaldehyde and stored in 4 degrees Celsius for 8 h. Following that tissues were submerged in 15% and 30% sucrose for 24 h each. Fixed tissues were trimmed and placed in OCT (Sakura Finetek) filled cryomoulds (Sakura Finetek) and flash frozen before long term storage in -80°C.

For flow cytometry, wounds were collected in 500 μl of serum free media, to which Liberase (Merck) was added, final concentration 200 μg/mL and placed at 37°C for 1 h, with occasional agitation to dissociated host cells. Equal volume FACS staining buffer (FCSB, PBS plus 2% heat-inactivated fetal bovine serum [FBS, Gibco] and 0.2 mM EDTA [Gibco]) was then added to stop the reaction. Cells were strained using a 40 um cell strainer (SPL Life Sciences), centrifuged and resuspended in 100μl FCSB). Following counting on a Countess 3 cell counter (Thermofisher), ∼5 x 10^6^ cells were taken for further processing and examination via flow cytometry.

### Murine neutrophil extraction and culture

Murine bone marrow neutrophils were obtained as previously described [71]. Briefly, femurs and tibias were removed from 6-9 week old female C57BL/6 mice, placed on ice and flushed with cold human plasma like media (HPLM, Gibco) using a 25G needle (Terumo). Cells were resuspended in MACs buffer (PBS [Gibco] plus 0.5% FBS, 2 mM EDTA) and run through a 40 um cell strainer before being isolated via negative selection using a Mojosort mouse neutrophil isolation kit (Biolegend). Cells were first incubated for 15 min on ice with a biotin-labelled antibody cocktail, followed incubation with streptavidin-conjugated magnetic beads for another 15 min on ice. The washed resuspension was then added to an LS column (Miltenyi Biotec) and placed on a magnetic separator (Miltenyi Biotech). The flow through (containing purified neutrophils was collected, ∼6-8M/mouse). For the majority of subsequent experimentation, cells were resuspended in HPLM plus 10% heat-inactivated FBS and plated into 24 well plates that has been precoated with 0.0001% poly-L-lysine (Sigma-Aldrich), 500k cells/ 1 mL/ well. For immunofluorescence, cells were first resuspended in Opti-Mem (Gibco), plated onto 12 mm glass coverslips (Fisher Scientific)(also precoated with poly-L-lysine) and stained with cytoplasm dye CellTracker Blue (Thermofisher) final concentration 5 μM for 30 min, before media was removed and replaced with fresh HPLM.

### Human neutrophil extraction and culture

Primary human neutrophils were obtained from human whole blood using negative selection with the MACSxpress Whole Blood Neutrophil Isolation Kit (Miltenyi Biotec), following manufacturers instructions. Briefly, 8 mL of whole blood (freshly collected into EDTA-coated tubes at Fullerton Health, NTU) was combined with an antibody cocktail and buffer solution. After thorough mixing this was placed on a strong magnet (Miltenyi Biotec), and left for 15 min. All antibody/magnetic bead coated cells then coagulate, leaving residual supernatant containing neutrophils alone, which was removed and resuspended in PBS for counting. Yield was ∼16-24M cells. Neutrophil purity was determined using antibodies against human CD45 and CD15 (2D1, HI98, Thermofisher).

### Immunofluorescence microscopy

For in vitro neutrophils on coverslips – at the given timepoints, 750 μl of media was removed from each well and replaced with 250 μl 8% PFA (final concentration 4%) and fixed for 15 min at 37°C. After thrice washing with PBS, cells were permeabilised with 0.1% Triton for 5 min), then blocked for 1 h with 1% bovine serum albumin (BSA, Sigma-Alridich). For cryosections on glass slides – Sections were first incubated in IHC Antigen Retrieval Solution (Invitrogen) for 20 min at 60°C, before being washed thrice with Milli-Q (Elgar Labwater) and once with PBS. Containing cells were permeabilise by dipping into 0.5% Triton in PBS for 5 min, followed by thrice washing in PBS. Sections were then blocked using blocking serum (PBS plus 5% FBS, 5% BSA, 0.05% Triton) for 1 h. Both were then incubated with polyclonal rabbit antibodies against Anti-Streptococcus Group D antigen (1:500 dilution from whole serum aliquot)(#12-6231D, American Research Products), in either 1% BSA (coverslips) or antibody diluent solution (PBS plus 1% FBS, 1% BSA, 0.05% Triton) overnight at 4°C. After thrice washing with PBS, were then incubated with goat anti-rabbit Alexa-488 1:1000 (A11034, Invitrogen), in same respectively diluents for 1 h at RT. Samples were again washed thrice again with PBS, with DNA then stained with 400uM propidium iodide (Invitrogen) (coverslips) for 15 min, or 10 ug/mL DAPI (Thermofisher) for 5 min. Samples were then mounted using ProLong^TM^ Glass Antifade Mountant (Thermofisher) and imaged at 63x magnification on a Carl Zeiss LSM 780 laser scanning confocal microscope (NTU Optical Bio-Imaging Centre [NOBIC] imaging facility, SCELSE).

### Flow cytometry

Flow cytometry was performed as described previously [12]. Briefly, excised skin samples were placed in 1.5 mL of liberase (2.5 U/ml) prepared in cell culture media. The mixture was incubated with agitation for 1 h at 37°C with 5% CO_2_. Dissociated cells were then passed through a 70 μm cell strainer and spun down at 1350 rpm for 5 min at 4°C. The enzymatic solution was then aspirated, and cells blocked in 500 μl of fluorescence-activated cell sorting (FACS) buffer [2% FBS and 0.2 mM EDTA in PBS (Gibco; Thermo Fisher Scientific)]. Cells (^∼^10^7^ per sample) were then incubated with 10 μl of Fc-blocker (anti-CD16/CD32 antibody; BioLegend) for 30 min, followed by incubation with an anti-mouse CD45, CD11b, and Ly6G (neutrophils), or CD45, CD11b, and F4/80 (macrophages) (BioLegend) (All 1:100 dilution) for 30 min. Cells were again washed in FACS buffer before being fixed in 4% PFA for 15 min at 4°C. Cells were washed a final time then resuspended and analysed using the BD LSRFortessa X-20 Cell Analyzer (Becton Dickinson). Compensation was performed using the AbC Total Antibody Compensation Bead Kit (Thermo Fisher Scientific) as per the manufacturer’s instructions.

### Lactate dehydrogenase (LDH) assay

Neutrophil death was determined using a cytotoxicity detection kit (Merck) following manufacturers instructions, with modifications to determine both approximate cell death from cell supernatants as well as remaining cell lysates to overcome potential bacterial interference with LDH readings [78]. Briefly, alongside experimental condition wells an identical well of cells was lysed with the addition of 10% Triton (final concentration 0.1%)(Sigma-Aldrich) 30 min prior to the experimental endpoint. This well was then flushed and the supernatant used as the ‘100% Triton kill’ well to which all other cell death is determined in proportion to.

Following initial sampling of supernatants, all media was removed from each well and replaced with 1 mL of HPLM + 0.1% Triton for 15 min (i.e. to lyse all remaining viable cells). This was then sampled and run alongside initial supernatant samples. After centrifugation, 50 ul of sample was combined with 50 μl of freshly prepared reagent and left for 30 min, after which the plate was read at 490nm in a Tecan Infinite 200 PRO spectrophotometer. NB for 24 h assay readings, gentamicin was added directly to the well (final concentration 200ug/mL) at 6 h p.i.

### Bacterial clearance assay

Following OD normalisation, *E. faecalis* bacteria were incubated with 10% mouse serum (Opsonised) or 10% heat-inactivated FBS (Non-opsonised) for 15 min at 37°C, after which were washed and resuspended in PBS and used for infection. Neutrophils were infected with MOI 1 or MOI 10 of bacteria for 2 or 6 h, after which the supernatants were sampled (and serially diluted on BHI agar for CFU enumeration of extracellular bacteria). All infectious media was then removed, and replaced with 1 mL of HPLM + 200 ug/mL gentamicin for 30 min. Media was then removed and neutrophils washed carefully twice with PBS, before being lysed with 1 mL PBS plus 0.1% Triton. This solution was then also serially diluted as above for CFU enumeration of intracellular bacteria. For the 24 h timepoint, at 6 h media was removed and replaced with HPLM plus 200 μg/mL gentamicin for 1 h, after which it was removed at replaced with HPLM plus 100 μg/mL gentamicin for the remainder of the experiment. To confirm the efficacy of gentamicin bacterial killing, *E. faecalis* bacteria were cultured in HPLM media alone for 6 h, followed by the addition of 200 μg/mL gentamicin. At 7 h this was left as in, or the media removed and replaced with 100 μg/mL gentamicin in media. Supernatant was removed and serially diluted from all wells at 6, 7, and 24 h, and plated out on BHI agar for CFU enumeration.

### Transmission Electron microscopy

Cultures of neutrophils with and without bacteria were fixed for 1 h in 2.5% glutaraldehyde (Ted Pella, Inc.) in PBS, pH 7.3, washed in buffer, post-fixed for 1 h in 1% osmium tetroxide with 1.5% Potassium Ferrocyanide in PBS, then dehydrated in an ethanol series, absolute acetone and embedded in Araldite 502 resin (Ted Pella, Inc). Ultra-sections were cut at 100 nm on a Leica EM UC6 Ultramicrotome, collected on 200 mesh copper grids covered with a Formvar carbon support film (Electron Microscopy Sciences), and stained for 8 min in lead citrate stain. Photographs were taken with Transmission Electron Microscopy (JEM-1400Flash, JEOL Ltd.) at 100kV with a bottom mounted high-sensitivity sCMOS camera.

### Multiplex cytokine analysis

Following infection of murine neutrophils with MOI 1 of *E. faecalis* OG1RF above, or stimulation with 10 μg/mL *S. pyogenes* lipoteichoic acid (Sigma Aldrich), cell culture supernatants were taken at 6 h and 24 h p.i. These were prepared as per manufacturer’s instructions and loaded onto a Bio-Plex PRO^TM^ Mouse Cytokine 23-plex Assay (Bio-Rad, California) to evaluating total cytokine concentrations. Sample processing, plate analysis and calculation of cytokine concentrations were performed at Bio-Rad Laboratories, Singapore with the assistance of their technical team.

### Ethics statement

All animal procedures were reviewed and approved by Institutional Animal Care and Use Committee (IACUC) in Nanyang Technological University (AUP# A19061) and performed accordingly. All procedures involving human neutrophils has been approved by the NTU Institutional Review Board, Reference Number: IRB-2020-06-005.

### Statistical analysis

Statistical analyses were performed utilising GraphPad Prism software (Version 10). Figure legends list specific test employed. For data sets involving 2 groups, a unpaired t-test was used. For datasets involving 3 or more groups a One-Way ANOVA with multiple comparisons test was performed. Details on specific tests applied and data obtained are contained within Figure legends.

## Acknowledgements

This work was supported by the National Research Foundation and Ministry of Education Singapore under its Research Centre of Excellence Programme (to the Singapore Centre for Environmental Life Sciences Engineering, SCELSE), and by the Singapore Ministry of Education under its Tier 2 program (MOE2019-T2-2-089) awarded to K.A.K. C.J.S was supported by SCELSE Seed Funding (SF-06). R.A.G.d.S. was supported by the National Research Foundation, Prime Minister’s Office, Singapore, under its Campus for Research Excellence and Technological Enterprise (CREATE) program, through core funding of the Singapore-MIT Alliance for Research and Technology (SMART) Antimicrobial Resistance Interdisciplinary Research Group (AMR IRG).

The authors thank the Electron Microscopy Unit (https://medicine.nus.edu.sg/core-facilities/electron-microscopy-unit-emu/) in National University of Singapore and the team, particularly Low Kay En, for their support and assistance in the TEM work.

## Author contributions

Conceptualisation, CJS, SLW, KAK; Methodology, CJS; Investigation, CJS, RAGS, NSOJ, HA; Writing – original draft, CJS; Writing – review and editing, CJS, HA, SLW, KAK; Resources, KAK; Funding acquisition, SLW and KAK

## References

1. Clinton, A. and T. Carter, Chronic Wound Biofilms: Pathogenesis and Potential Therapies. Laboratory Medicine, 2015. 46(4): p. 277–284.

2. Martinengo, L., et al., Prevalence of chronic wounds in the general population: systematic review and meta-analysis of observational studies. Annals of Epidemiology, 2019. 29: p. 8–15.

3. Mottola, C., et al., Polymicrobial biofilms by diabetic foot clinical isolates. Folia Microbiol (Praha), 2016. 61(1): p. 35–43.

4. Thanganadar Appapalam, S., A. Muniyan, K. Vasanthi Mohan, and R. Panchamoorthy, A Study on Isolation, Characterization, and Exploration of Multiantibiotic-Resistant Bacteria in the Wound Site of Diabetic Foot Ulcer Patients. Int J Low Extrem Wounds, 2019: p. 1534734619884430.

5. Małecki, R., K. Klimas, and A. Kujawa, Different Patterns of Bacterial Species and Antibiotic Susceptibility in Diabetic Foot Syndrome with and without Coexistent Ischemia. J Diabetes Res, 2021. 2021: p. 9947233.

6. Fisher, K. and C. Phillips, The ecology, epidemiology and virulence of Enterococcus. Microbiology (Reading), 2009. 155(Pt 6): p. 1749–1757.

7. Kao, P.H.N. and K.A. Kline, Dr. Jekyll and Mr. Hide: How Enterococcus faecalis Subverts the Host Immune Response to Cause Infection. Journal of Molecular Biology, 2019. 431(16): p. 2932–2945.

8. Arias, C.A. and B.E. Murray, The rise of the Enterococcus: beyond vancomycin resistance. Nat Rev Microbiol, 2012. 10(4): p. 266–78.

9. Ch’ng, J.H., et al., Biofilm-associated infection by enterococci. Nat Rev Microbiol, 2019. 17(2): p. 82–94.

10. Bertuccini, L., M.G. Ammendolia, F. Superti, and L. Baldassarri, Invasion of HeLa cells by Enterococcus faecalis clinical isolates. Med Microbiol Immunol, 2002. 191(1): p. 25–31.

11. Zou, J. and N. Shankar, The opportunistic pathogen Enterococcus faecalis resists phagosome acidification and autophagy to promote intracellular survival in macrophages. Cell Microbiol, 2016. 18(6): p. 831–43.

12. da Silva, R.A.G., et al., Enterococcus faecalis alters endo-lysosomal trafficking to replicate and persist within mammalian cells. PLOS Pathogens, 2022. 18(4): p. e1010434.

13. Archambaud, C., et al., Enterococcus faecalis: an overlooked cell invader. Microbiol Mol Biol Rev, 2024. 88(3): p. e0006924.

14. Burn, G.L., et al., The Neutrophil. Immunity, 2021. 54(7): p. 1377–1391.

15. Watson, R.W., et al., Neutrophils undergo apoptosis following ingestion of Escherichia coli. J Immunol, 1996. 156(10): p. 3986–92.

16. Rotstein, D., J. Parodo, R. Taneja, and J.C. Marshall, Phagocytosis of Candida albicans induces apoptosis of human neutrophils. Shock, 2000. 14(3): p. 278–83.

17. Kobayashi, S.D., et al., Bacterial pathogens modulate an apoptosis differentiation program in human neutrophils. Proc Natl Acad Sci U S A, 2003. 100(19): p. 10948–53.

18. Kobayashi, S.D., N. Malachowa, and F.R. DeLeo, Influence of Microbes on Neutrophil Life and Death. Frontiers in cellular and infection microbiology, 2017. 7: p. 159–159.

19. van Zandbergen, G., et al., Chlamydia pneumoniae multiply in neutrophil granulocytes and delay their spontaneous apoptosis. J Immunol, 2004. 172(3): p. 1768–76.

20. Aga, E., et al., Inhibition of the spontaneous apoptosis of neutrophil granulocytes by the intracellular parasite Leishmania major. J Immunol, 2002. 169(2): p. 898–905.

21. Chen, A. and H.S. Seifert, Neisseria gonorrhoeae-mediated inhibition of apoptotic signalling in polymorphonuclear leukocytes. Infection and immunity, 2011. 79(11): p. 4447–4458.

22. McCracken, J.M., L.C. Kinkead, R.L. McCaffrey, and L.A. Allen, Francisella tularensis Modulates a Distinct Subset of Regulatory Factors and Sustains Mitochondrial Integrity to Impair Human Neutrophil Apoptosis. J Innate Immun, 2016. 8(3): p. 299–313.

23. Guzmàn, C.A., C. Pruzzo, and L. Calegari, Enterococcus faecalis: specific and non-specific interactions with human polymorphonuclear leukocytes. FEMS Microbiol Lett, 1991. 68(2): p. 157–62.

24. Vanek, N.N., et al., Enterococcus faecalis aggregation substance promotes opsonin-independent binding to human neutrophils via a complement receptor type 3-mediated mechanism. FEMS Immunology & Medical Microbiology, 1999. 26(1): p. 49–60.

25. Gaglani, M.J., C.J. Baker, and M.S. Edwards, Contribution of antibody to neutrophil-mediated killing of Enterococcus faecalis. J Clin Immunol, 1997. 17(6): p. 478–84.

26. Harvey, B.S., C.J. Baker, and M.S. Edwards, Contributions of complement and immunoglobulin to neutrophil-mediated killing of enterococci. Infection and immunity, 1992. 60(9): p. 3635–3640.

27. Arduino, R.C., B.E. Murray, and R.M. Rakita, Roles of antibodies and complement in phagocytic killing of enterococci. Infect Immun, 1994. 62(3): p. 987–93.

28. Rakita, R.M., et al., Enterococcus faecalis bearing aggregation substance is resistant to killing by human neutrophils despite phagocytosis and neutrophil activation. Infect Immun, 1999. 67(11): p. 6067–75.

29. Süssmuth, S.D., et al., Aggregation substance promotes adherence, phagocytosis, and intracellular survival of Enterococcus faecalis within human macrophages and suppresses respiratory burst. Infect Immun, 2000. 68(9): p. 4900–6.

30. Chong, K.K.L., et al., Enterococcus faecalis Modulates Immune Activation and Slows Healing During Wound Infection. The Journal of infectious diseases, 2017. 216(12): p. 1644–1654.

31. Kim, M.H., et al., Dynamics of neutrophil infiltration during cutaneous wound healing and infection using fluorescence imaging. J Invest Dermatol, 2008. 128(7): p. 1812–20.

32. Kanno, E., et al., Wound healing in skin promoted by inoculation with Pseudomonas aeruginosa PAO1: The critical role of tumor necrosis factor-α secreted from infiltrating neutrophils. Wound Repair and Regeneration, 2011. 19(5): p. 608–621.

33. Hallinen, K.M., K.A. Guardiola-Flores, and K.B. Wood, Fluorescent reporter plasmids for single-cell and bulk-level composition assays in E. faecalis. PLOS ONE, 2020. 15(5): p. e0232539.

34. Zurek, O.W., K.B. Pallister, and J.M. Voyich, Staphylococcus aureus Inhibits Neutrophil-derived IL-8 to Promote Cell Death. J Infect Dis, 2015. 212(6): p. 934–8.

35. Van den Bossche, S., et al., Bacterial Interference With Lactate Dehydrogenase Assay Leads to an Underestimation of Cytotoxicity. Frontiers in Cellular and Infection Microbiology, 2020. 10(494).

36. Paulsen, I.T., et al., Role of mobile DNA in the evolution of vancomycin-resistant Enterococcus faecalis. Science, 2003. 299(5615): p. 2071–4.

37. Ch’ng, J.H., et al., Heme cross-feeding can augment Staphylococcus aureus and Enterococcus faecalis dual species biofilms. Isme j, 2022.

38. Hsu, Y.P., et al., Full color palette of fluorescent d-amino acids for in situ labeling of bacterial cell walls. Chem Sci, 2017. 8(9): p. 6313–6321.

39. Mitchell, G., C. Chen, and A. Portnoy Daniel, Strategies Used by Bacteria to Grow in Macrophages. Microbiology Spectrum, 2016. 4(3): p. 10.1128/microbiolspec.mchd-0012-2015.

40. Kobayashi, S.D., N. Malachowa, and F.R. DeLeo, Neutrophils and Bacterial Immune Evasion. J Innate Immun, 2018. 10(5-6): p. 432–441.

41. Kennedy, A.D. and F.R. DeLeo, Neutrophil apoptosis and the resolution of infection. Immunologic Research, 2009. 43(1): p. 25–61.

42. Kobayashi, S.D., et al., Rapid neutrophil destruction following phagocytosis of Staphylococcus aureus. J Innate Immun, 2010. 2(6): p. 560–75.

43. Simons, M.P., W.M. Nauseef, T.S. Griffith, and M.A. Apicella, Neisseria gonorrhoeae delays the onset of apoptosis in polymorphonuclear leukocytes. Cell Microbiol, 2006. 8(11): p. 1780–90.

44. McCaffrey, R.L. and L.A. Allen, Francisella tularensis LVS evades killing by human neutrophils via inhibition of the respiratory burst and phagosome escape. J Leukoc Biol, 2006. 80(6): p. 1224–30.

45. van Zandbergen, G., et al., Cutting edge: neutrophil granulocyte serves as a vector for Leishmania entry into macrophages. J Immunol, 2004. 173(11): p. 6521–5.

46. Yoshiie, K., H.Y. Kim, J. Mott, and Y. Rikihisa, Intracellular infection by the human granulocytic ehrlichiosis agent inhibits human neutrophil apoptosis. Infection and immunity, 2000. 68(3): p. 1125–1133.

47. Nazareth, H., S.A. Genagon, and T.A. Russo, Extraintestinal pathogenic Escherichia coli survives within neutrophils. Infect Immun, 2007. 75(6): p. 2776–85.

48. Gresham, H.D., et al., Survival of Staphylococcus aureus inside neutrophils contributes to infection. J Immunol, 2000. 164(7): p. 3713–22.

49. Millán, D., C. Chiriboga, M.A. Patarroyo, and M.R. Fontanilla, Enterococcus faecalis internalization in human umbilical vein endothelial cells (HUVEC). Microbial Pathogenesis, 2013. 57: p. 62–69.

50. Horsley, H., et al., Enterococcus faecalis Subverts and Invades the Host Urothelium in Patients with Chronic Urinary Tract Infection. PLOS ONE, 2013. 8(12): p. e83637.

51. Horsley, H., D. Dharmasena, J. Malone-Lee, and J.L. Rohn, A urine-dependent human urothelial organoid offers a potential alternative to rodent models of infection. Scientific Reports, 2018. 8(1): p. 1238.

52. Gentry-Weeks Claudia, R., et al., Survival of Enterococcus faecalis in Mouse Peritoneal Macrophages. Infection and Immunity, 1999. 67(5): p. 2160–2165.

53. Zou, J. and N. Shankar, Enterococcus faecalis Infection Activates Phosphatidylinositol 3-Kinase Signaling To Block Apoptotic Cell Death in Macrophages. Infection and Immunity, 2014. 82(12): p. 5132.

54. Nunez, N., et al., The unforeseen intracellular lifestyle of Enterococcus faecalis in hepatocytes. Gut Microbes, 2022. 14(1): p. 2058851.

55. Hertzén, E., et al., Intracellular Streptococcus pyogenes in Human Macrophages Display an Altered Gene Expression Profile. PLOS ONE, 2012. 7(4): p. e35218.

56. Celik, C., et al., Decoding the complexity of delayed wound healing following Enterococcus faecalis infection. Elife, 2024. 13.

57. Chaves, M.M., et al., The role of dermis resident macrophages and their interaction with neutrophils in the early establishment of Leishmania major infection transmitted by sand fly bite. PLoS Pathog, 2020. 16(11): p. e1008674.

58. Stocks, C.J., et al., Frontline Science: LPS-inducible SLC30A1 drives human macrophage-mediated zinc toxicity against intracellular Escherichia coli. J Leukoc Biol, 2021. 109(2): p. 287–297.

59. Lotz, S., et al., Highly purified lipoteichoic acid activates neutrophil granulocytes and delays their spontaneous apoptosis via CD14 and TLR2. J Leukoc Biol, 2004. 75(3): p. 467–77.

60. Colotta, F., et al., Modulation of granulocyte survival and programmed cell death by cytokines and bacterial products. Blood, 1992. 80(8): p. 2012–20.

61. Adachi, S., et al., In vivo administration of granulocyte colony-stimulating factor promotes neutrophil survival in vitro. Eur J Haematol, 1994. 53(3): p. 129–34.

62. Begley, C.G., et al., Purified colony-stimulating factors enhance the survival of human neutrophils and eosinophils in vitro: a rapid and sensitive microassay for colony-stimulating factors. Blood, 1986. 68(1): p. 162–6.

63. Saba, S., G. Soong, S. Greenberg, and A. Prince, Bacterial stimulation of epithelial G-CSF and GM-CSF expression promotes PMN survival in CF airways. Am J Respir Cell Mol Biol, 2002. 27(5): p. 561–7.

64. Choi, M., et al., Inhibition of NF-kappaB by a TAT-NEMO-binding domain peptide accelerates constitutive apoptosis and abrogates LPS-delayed neutrophil apoptosis. Blood, 2003. 102(6): p. 2259–67.

65. Roilides, E., T.J. Walsh, P.A. Pizzo, and M. Rubin, Granulocyte colony-stimulating factor enhances the phagocytic and bactericidal activity of normal and defective human neutrophils. J Infect Dis, 1991. 163(3): p. 579–83.

66. Salamone, G., et al., Promotion of neutrophil apoptosis by TNF-alpha. J Immunol, 2001. 166(5): p. 3476–83.

67. Murray, J., et al., Regulation of Neutrophil Apoptosis by Tumor Necrosis Factor-α: Requirement for TNFR55 and TNFR75 for Induction of Apoptosis In Vitro. Blood, 1997. 90(7): p. 2772–2783.

68. Tien, B.Y.Q., et al., Enterococcus faecalis Promotes Innate Immune Suppression and Polymicrobial Catheter-Associated Urinary Tract Infection. Infect Immun, 2017. 85(12).

69. McCaffrey, R.L., et al., Multiple mechanisms of NADPH oxidase inhibition by type A and type B Francisella tularensis. J Leukoc Biol, 2010. 88(4): p. 791–805.

70. Schwartz, J.T., et al., Francisella tularensis inhibits the intrinsic and extrinsic pathways to delay constitutive apoptosis and prolong human neutrophil lifespan. J Immunol, 2012. 188(7): p. 3351–63.

71. Kao, P.H.-N., et al., Enterococcus faecalis suppresses Staphylococcus aureus-induced NETosis and promotes bacterial survival in polymicrobial infections. FEMS Microbes, 2023. 4: p. xtad019.

72. Carneiro, B.A. and W.S. El-Deiry, Targeting apoptosis in cancer therapy. Nature Reviews Clinical Oncology, 2020. 17(7): p. 395–417.

73. Ebert, G., et al., Eliminating hepatitis B by antagonizing cellular inhibitors of apoptosis. Proc Natl Acad Sci U S A, 2015. 112(18): p. 5803–8.

74. Speir, M., et al., Eliminating Legionella by inhibiting BCL-XL to induce macrophage apoptosis. Nature Microbiology, 2016. 1(3): p. 15034.

75. Antypas, H., et al., Fsr quorum sensing system restricts biofilm growth and activates inflammation in enterococcal infective endocarditis. bioRxiv, 2025.

76. Dunny, G.M., R.A. Craig, R.L. Carron, and D.B. Clewell, Plasmid transfer in Streptococcus faecalis: Production of multiple sex pheromones by recipients. Plasmid, 1979. 2(3): p. 454–465.

77. King, M.D., et al., Emergence of community-acquired methicillin-resistant Staphylococcus aureus USA 300 clone as the predominant cause of skin and soft-tissue infections. Ann Intern Med, 2006. 144(5): p. 309–17.

78. Van den Bossche, S., et al., Bacterial Interference With Lactate Dehydrogenase Assay Leads to an Underestimation of Cytotoxicity. Front Cell Infect Microbiol, 2020. 10: p. 494.

